# ConvMut: Exploration of viral convergent mutations along phylogenies

**DOI:** 10.1101/2024.12.16.628620

**Authors:** Tommaso Alfonsi, Anna Bernasconi, Emma Fanfoni, Cesare Ernesto Maria Gruber, Fabrizio Maggi, Daniele Focosi

**Affiliations:** Department of Electronics, Information, and Bioengineering. Politecnico di Milano, Milan, Italy; Laboratory of Virology and Laboratories of Biosecurity, National Institute for Infectious Diseases Lazzaro Spallanzani. IRCCS, Rome, Italy; Bioinformatics Research Unit in Infectious Diseases, National Institute for Infectious Diseases Lazzaro Spallanzani. IRCCS, Rome, Italy; North-Western Tuscany Blood Bank. Pisa University Hospital, Pisa, Italy

## Abstract

Convergent evolution in protein antigens is common across pathogens and has also been documented in SARS-CoV-2 (hCoV-19); the most likely reason is the need to evade the selective pressure exerted by previous infection- or vaccine-elicited immunity. There is a pressing need for tools that allow automated analysis of convergent mutations.

In response to this need, we developed ConvMut, a tool to analyze genetic sequence data to identify patterns of recurrent mutations in SARS-CoV-2 evolution. To this end, we exploited the granular phylogenetic tree representation developed by PANGO, allowing us to observe what we call deltas, i.e., groups of mutations that are acquired on top of the immediately upstream tree nodes. Deltas comprise amino acid substitutions, insertions, and deletions. ConvMut can perform individual protein analysis to identify the most common single mutations acquired independently in a given subtree (starting from a user-selected root). Such mutations are represented in a barplot that can be sorted by frequency or position, and filtered by region of interest. Lineages are then gathered into clusters according to their sets of shared mutations. Finally, an interactive graph orders the evolutionary steps of clusters, details the acquired amino acid changes for each sublineage, and allows us to trace the evolutionary path until a selected lineage.

Other unique tools are paired with the main functionality of ConvMut to support a complete analysis, such as a frequency analysis for a given nucleotide or amino acid changes at a given residue across a selected phylogenetic subtree.

ConvMut will facilitate the design of antiviral anti-Spike monoclonal antibodies and Spike-based vaccines with longer-lasting efficacy, minimizing development and marketing failures.

## Introduction

SARS-CoV-2 represents a panzootic of placental mammals [1], with multiple examples of reverse zoonosis documented so far [2,3,4] and the persistence of variants nearly extinct in humans [5,6]. Such massive circulation across different species creates 28.4 substitutions per year and remains about 10-fold faster than for other RNA viruses established in humans since much more time (e.g., measles, mumps, RSV, dengue, Zika, WNV, enterovirus D68, or influenza virus A/H3N2, as documented by unprecedented genome sequencing efforts throughout the world with almost 17.5 million sequences that have been shared via the GISAID data science initiative [7]. From December 2019 to September 2025, more than 5,300 sublineages have been annotated by GISAID following PANGO designations [8] and more than 3,000 have been designated since December 2021, falling under the WHO umbrella of the Omicron variant of concern (VoC) [9]. Such a granular phylogenetic tree represents a unique opportunity for studying convergent evolution. Globally, research has confirmed trends toward greater serological distance [10], translating into higher immune escape and finally into progressively higher reproductive numbers [11]. In proteomics, convergent evolution refers to the independent acquisition of genetic mutations leading to identical amino acid changes. The aim of those changes is adaptation to environmental variables, which for a pathogen implies immune evasion.

Convergent evolution in SARS-CoV-2 has been reviewed in detail elsewhere [12,13] and has led, e.g., to sudden failures of most of the therapeutic anti-Spike monoclonal antibodies [14]. To date, most insights on SARS-CoV-2 convergent evolution stem from manual work run by volunteers on social platforms, and no automated tool has been developed yet. We present here ConvMut, a comprehensive software for the analysis of convergent evolution in SARS-CoV-2 proteins.

## Materials and Methods

In the following sections, we provide a set of useful definitions and outline the data processing pipeline.

### Concepts definition

We keep note of the PANGO lineage nomenclature, updated regularly [8]; thanks to the hierarchical nature of Pango nomenclature, we can completely reconstruct the implied hierarchy, focusing on the alias naming scheme that represents the full hierarchical provenance. For instance, if JN is an alias for B.1.1.529.2.86.1, then JN.1 is represented by the *unaliased* lineage B.1.1.529.2.86.1.1 in our hierarchical data structure.

For each lineage, its *constellation* is a list of all the nucleotide mutations (expressed with regard to the reference genome) that are progressively acquired by one tree-node (either representing a designated or undesignated lineage) with respect to its previous node in the hierarchy. This traces a path along the tree, starting from its root to the lineage represented in that line. Different tree levels are marked with the symbol “>“; possibly, multiple substitutions are incorporated in the same level (meaning that these are acquired jointly by the observed node). As an example, the lineage B.1.617.2 can be associated with the path “C14408T > C241N > C3037T > A23403G > G29742N > T22917G > T27638C > G28881T > G210N > C21618G > G29402T > C23604G > C25469T > C22995A, C27752T > A28461G”, indicating that –starting from the root lineage designated as B– 15 levels of the phylogeny have been traversed. At each step on the tree, one mutation is acquired with respect to the previous node (first C14408T, then C241N, etc.). Finally, the two mutations C22995A, C27752T are acquired by the parent node of B.1.617.2 with respect to the grandparent node, whereas A28461G is acquired by B.1.617.2 with respect to its parent node.

### Data input pipeline

#### GISAID

The ConvMut application is available on GISAID EpiCoV [7], where it leverages a file (completely computed by GISAID internal pipelines) that contains lines composed of Lineage, Constellation, the earliest collection date of the lineage, and lineage sequence count. The mutation calling procedure is 1-based and employs the EPI_ISL_402124 (<u>WIV04</u>) sequence as the reference genome. The pipeline includes substitution, insertion, and deletion mutations. This data structure is updated daily and aligned with the full database of GISAID.

### Data processing pipeline

#### Tree reconstruction and translation

Once the input file has been prepared, we rebuild the original parent-child and child-parent links in the hierarchical phylogeny of SARS-CoV-2 by only including designated lineages.

We built *template genomes* for each lineage (by updating the reference genome with the alternative nucleotide indicated in the mutations that are assigned to the lineage). Note that reversions did not produce any change here.

For each template genome, we extract the open reading frames 1a, 1b, 3a, 6, 7a, 7b, 8, 9b, and the genes S, E, M, N, using common alignment tool MAFFT [15] and translate them into the corresponding proteins. After the alignment with the reference protein, we obtain a data structure mapping the mutated residues to the reference protein. The reference-aligned protein sequences are then useful to extract the wild type-lineage deltas, the inter-lineage deltas, and the mutated codons.

#### Inter-lineage deltas

For each considered lineage (*L*), we computed a *parent-delta*, i.e., the set of reference amino acids (with their position in the reference proteins) that have mutated, happening between *L* and *L*’s parent. Exceptionally, for those few lineages for which a parent was missing, we employed the first available ancestor in the dataset.

When *L* is recombinant, we computed a *delta* with respect to each of its parents. If an amino acid with its position was present in all deltas, it was inserted in the final *L*’s delta. Otherwise, we did not retain it (as we cannot rely on known breaking point coordinates for assigning a position to specific parents). We recognize that this leads to an approximation in the following cases:

1. in the observed position (*pos*), *LP1* exhibits *a1, LP2* exhibits *a2* and the recombinant lineage exhibits *a3*. Here, we will encode this situation as *a1_pos_a3*
2. in the observed position (*pos*), *LP1* exhibits *a1, LP2* exhibits *a2* and the recombinant lineage exhibits *a1*. Even if *a1* were to be considered an actual mutation (as derived from the portion of the genome inherited from *LP2*) this would not be recognized as a mutation, given that it is present in at least one of the parental lineages (i.e., *LP1*).

#### Wild type-lineage deltas

For each lineage, we computed a *wildtype-delta*, by comparing the amino acid exhibited in each position of the genome with the one present in the wild type. The “delta mutations” were stored in a dictionary. We noted that, for a given lineage, its parent-delta and wildtype-delta were not necessarily the same, as the following situations may arise:

- A certain mutation was included in parent-delta but not in wildtype-delta. This corresponds to a reversion, i.e., a mutation that re-instantiates the original wild-type genome.
- A certain mutation is included in the wildtype-delta but not in the parent-delta. Here, the observed lineage has inherited a mutation from its parent.

#### Quantifying a mutation convergence representativeness

We group lineages by mutations that are present in parent-deltas, thereby extracting counts of how many lineages have acquired that mutation with respect to their direct parent. These counts are used to draw a ranking among mutations; the *most convergent mutations* are those with the highest counts, as they are the most represented ones in parent-deltas (independently of the phylogenetic tree portion where this may happen).

Note that mutations in the same position contribute equally to computed counts, e.g., considering the position *pos*, if the *L1* lineage exhibits the mutation *a1_pos_a2*, and the *L2* lineage the mutation *a3_pos_a4*, the occurrence count for *pos* is increased by 2.

The previously illustrated approach may introduce a possible bias, as it ignores intermediate nodes that correspond to undesignated lineages. Specifically, all mutations acquired by a node in the tree that represents a non-designated lineage fall into the parent-delta of its designated offspring, thus possibly overestimating the mutation count.

## Web Application

ConvMut is implemented as an interactive Web data application based on the open-source Python framework Streamlit, deployed as a Docker-based container. It presents a landing page with documentation and three tabs, respectively, targeted to *convergence analysis, codon exploration*, and *mutation exploration*.

The application leverages pre-computed data structures. The corresponding workflow is run daily to capture potential updates in the input files. The data structures are then pre-loaded in the application when it is started, and then saved as session variables so that all tabs can reuse them and make the application more responsive.

In the following, for example purposes, we refer to ConvMut deployment on GISAID as of September 14th, 2025.

### Convergence analysis

This page allows us to analyze the most frequent convergent mutations in a specific SARS-CoV-2 protein within the entire phylogenetic tree or within a portion of the phylogeny starting from a user-selected *root* lineage. The interface user experience is built as a progressive workflow where the user is asked to select values to filter the analysis (unless default values are accepted). In the following, we define each phase.

#### Selection of a SARS-CoV-2 protein and root lineage

One *protein* among all the ones in SARS-CoV-2 must be selected. Depending on this choice, only positions/mutations on the selected protein are then considered. The default protein is ‘Spike’.

The *root* is the starting node of the subtree considered for the downstream analysis. Note that recombinant lineages are included in the analyzed subtree when at least one of their parent lineages is included in the subtree. ConvMut provides a tabular visualization of the subtree, where each line represents a *lineage* (plus its unaliased name), *parent-delta*, and *wildtype-delta*.

Additionally, ConvMut shows a horizontal barplot featuring the count of independent acquisitions of mutations occurring in each position of the selected protein along the selected subtree. All different mutation types occurring at a given position contribute to the global count for a given position (e.g., R346T, R346S, and S346I all contribute to the general count position 346). E.g., in a sub-tree rooted in JN.1.11.1, featuring 733 sub-lineages (plus JN.1.11.1 itself), we find 35 lineages whose parent-delta contains a mutation occurring on 346. Reversions count as regular mutations. The barplot can be sorted by position or number of occurrences.

#### Selection of positions of interest

The set of positions to be included in the following visualization is pre-filled by default using the positions exceeding a threshold set during the previous phase. By default, this is set to the value of the third most frequent position – should there be ties, at most 10 positions are included. Users can freely add or remove positions or change the entire set with an auto-complete component, e.g., when the analyzed protein is Spike, it is possible to restrict the analysis to specific regions, such as the receptor-binding domain (RBD: amino acid positions 319-541) or receptor-binding motif (RBM: amino acid positions 438-506).

We term *SP* the set of positions selected by the user; we compute and show 1) a series of pieplots representing the variability of different amino acid residues in each position in *SP*, and 2) a table including only the lineages within the selected subtree that include at least a position within *SP*. The table features the (aliased) lineage name, the unaliased lineage name, the cluster it belongs to, the mutations acquired by lineages in the cluster, and a string explaining how the cluster composition was computed.

For a meaningful, well-organized visualization of the map, we *cluster* lineages. Specifically – given a lineage *L*, its parent-delta, and wildtype-delta – *L*’s corresponding cluster is computed by taking only the positions in *SP* that are included in either *L*’s parent-delta or *L*’s wildtype-delta (or both). For instance, when *SP* = {346, 45}, *L1*’s parent-delta = {F456L, S680I}, and *L1*’s wildtype-delta = {S31-, R346S, F456L}, *L1*’s cluster will be characterized by {346,456}, then labelled c_346_456. Instead, with, *L2*’s parent-delta = {F456L}, and *L2*’s wildtype-delta = {Q480T}, *L2*’s cluster would be characterized by {456}, then labelled simply c_456.

Note that lineages grouped in the same *cluster* do not necessarily have the same mutations with respect to the user-selected root; instead, they must exhibit the same convergent mutations (even because of a reversion process).

#### Convergent mutations map

ConvMut finally plots a map to represent the complete convergent mutations’ landscape of the selected protein in the selected subtree, including only mutations corresponding to the positions chosen as described above. Nodes in the map are then a specific set of PANGO-designated lineages -termed *LV*- (determined as those whose parent-delta sets contain at least one mutation included in the positions of *SP*), and edges represent a parent-child (or ancestor-child) relationship between two lineages, labeled with mutation(s) contained in the *parent-delta* of the child node. Note that the graph is developed and visualized from left to right, by representing the root and older lineages in the left-most portion of the page and the leaves (most recent lineages) in the right-most portion.

We note that two nodes that are connected by an edge in the graph are not necessarily parent/child, as -given a child-it could happen that its direct parent is not contained in *LV* (and, in turn, its grandparent, etc.). Then, edges are drawn between a node and its closest PANGO-designated ancestor included in *LV*.

We also note that some clusters may enclose nodes whose incoming edges have different labels (either in the alternative amino acid residue or even in the position); this is consistent with the definition of a cluster, which gathers lineages that have the same mutations in their parent-delta or wildtype-delta, making the stage at which a mutation was acquired irrelevant.

### Codon exploration

This page allows us to explore mutations that occur in a specific codon of the selected protein. Starting from the amino acid residue, we provide information on the determinant codon. Users input a SARS-CoV-2 protein, a phylogenetic root (in the PANGO-lineage hierarchy) from which the analysis is produced, and a *position of interest* (among pre-filtered options only including those that are present in at least one *delta* in the selected sub-tree). Users can choose to visualize results based on codons or amino acid residues.

### Mutation exploration

This page allows users to investigate a given position on a selected protein, considering also its phylogenetic background. Consistent with the previous tab, we require a protein, root lineage, and position *P* to be inspected.

We identify with *LM* all the lineages in the considered subtree that include a mutation in the *P* position in their parent-delta. The user can visualize a tree-like representation of the initial lineages’ sub-tree, where we pruned all branches that do not present any lineage in *LM*; we also collapse tree levels not containing any lineage (or parent of a lineage) in *LM*. Mutation labels are colored to make differences in amino acids more apparent.

## Results

In this section, we report sample results for each of the ConvMut main functions.

Figure 1 shows intermediate results for the *Convergence analysis* when the following inputs are selected: `protein = S` (Spike)`, root = JN.1.11.1`, `threshold = 35` (to include positions of interest), and `selected positions` (from more represented to less represented) `= 346`, `31`, `572`.

**Figure 1.**
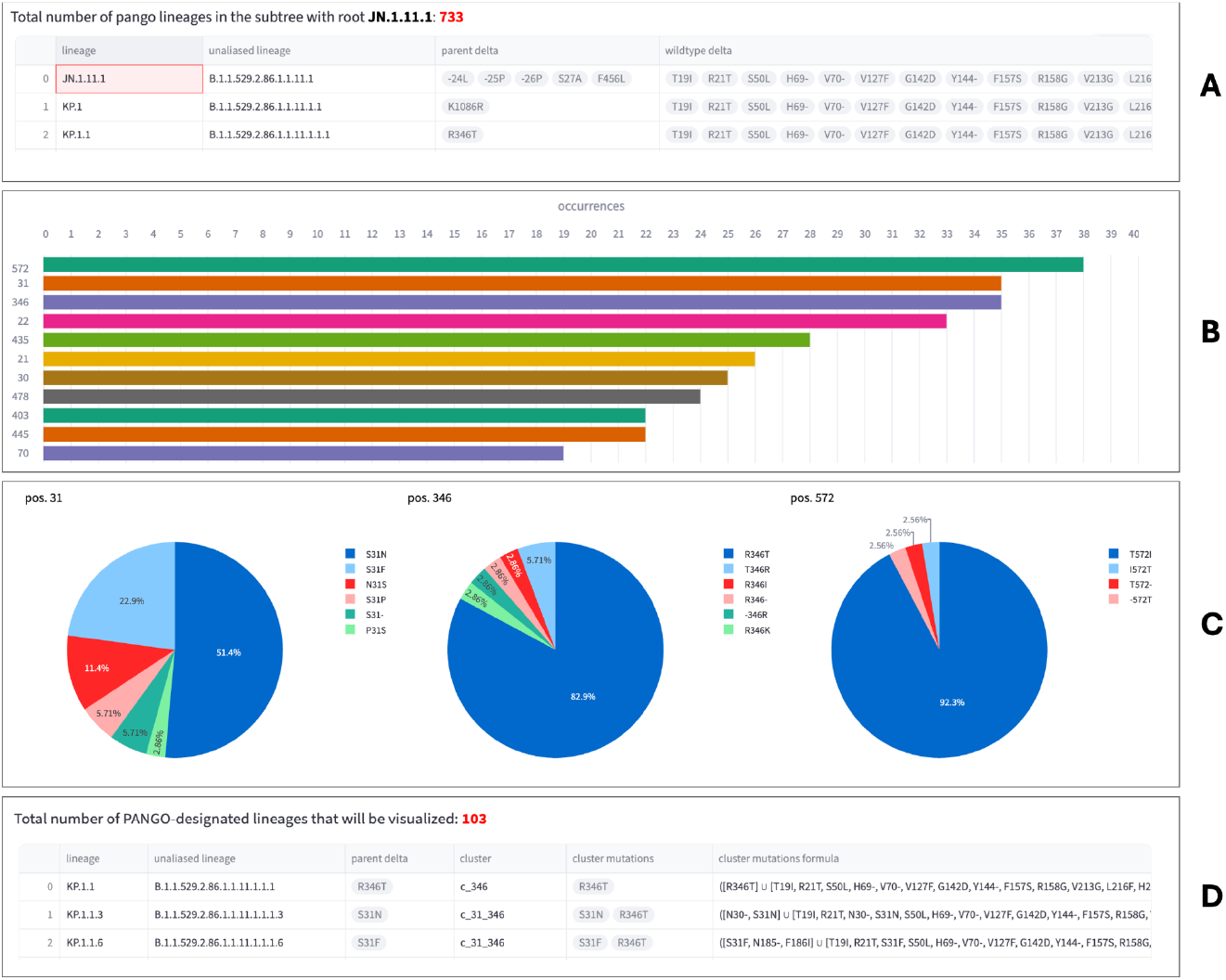
Intermediate results’ visualization. **(A)** Table with all the lineages that derive from the chosen root (JN.1.11.1, in the example); **(B)** Barplot that, for each position on the selected protein, counts how many lineages independently acquired a mutations in that position, sorted by count; **(C)** Pieplots that capture the variability of different amino acid residues in the three selected positions (i.e., those above the chosen threshold, 35, according to the barplot in panel B); **(D)** Table for detailing a lineage’s cluster name and its mutations for each of the lineages to be plotted in the graph.

Panel A illustrates a table with lineages (normal and unaliased) described by their parent-delta and wildtype-delta; Panel B shows the barplot whose y-axis represents positions along the chosen protein and the x-axis represents the count of lineages acquiring a mutation in that position; Panel C shows pieplots for each of the positions extracted above the set threshold (along the barplot shown in B), showing how mutations are distributed by alternative amino acid; Panel D presents a table of lineages to be visualized in the graph, detailing their cluster, the clusters’ specific mutations and how these were computed, according to the formula:

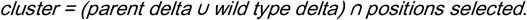

As a result of the previous parameters’ selection, a graph is rendered as shown in Figure 2.

**Figure 2.**
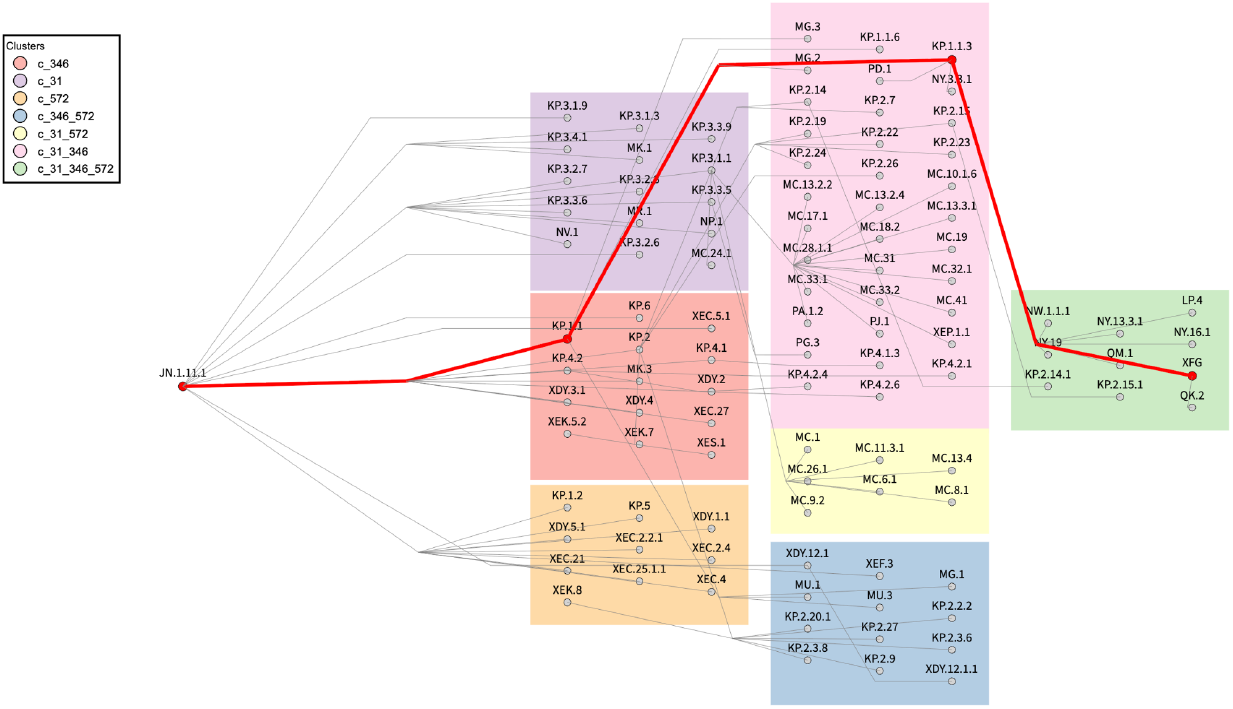
The ConvMut Convergence analysis tree, for Spike’s mutations `346, 31, 572` in the JN.1.11.1 subtree.

By selecting a specific lineage in the dropdown menu above the graph, we can highlight the specific relevant path of converging events on the subtree. For example, in Figure 2, we show the path of XFG, a recombinant lineage that is of particular interest at the time of writing [16]. While Figure 2 shows the visualized tree, Figure 3 zooms in on the observed path, obtained by exploiting the features of the implemented interactive visualization. Specifically, the chain to be followed from the root (left) to the selected lineage (right) can be described as follows: JN.1.11.1 acquired R346T in KP.1.1, which acquired S31N in KP.1.1.3. This, in turn, acquired T572I in NY.19, finally acquiring the triplet <P31S, T346R, I572T> in XFG.

**Figure 3.**
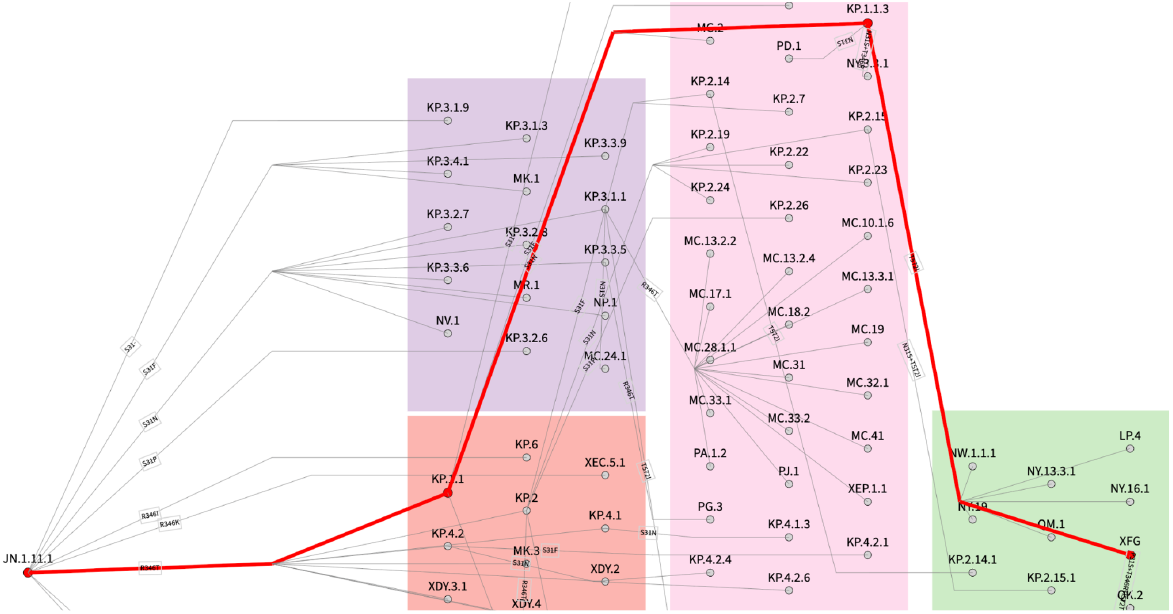
Zoomed visualization of the ConvMut Convergence analysis tree, for Spike’s mutations`, 346, 31, 572` in the JN.1.11.1 subtree. The path leading from the root JN.1.11.1 to the recent recombinant lineage XFG is highlighted in red.

Then, Figure 4 provides an overview of the possible output of the *Codon exploration* analysis, when the input `root = JN.1.11.1` and `pos = 346` are selected.

**Figure 4.**
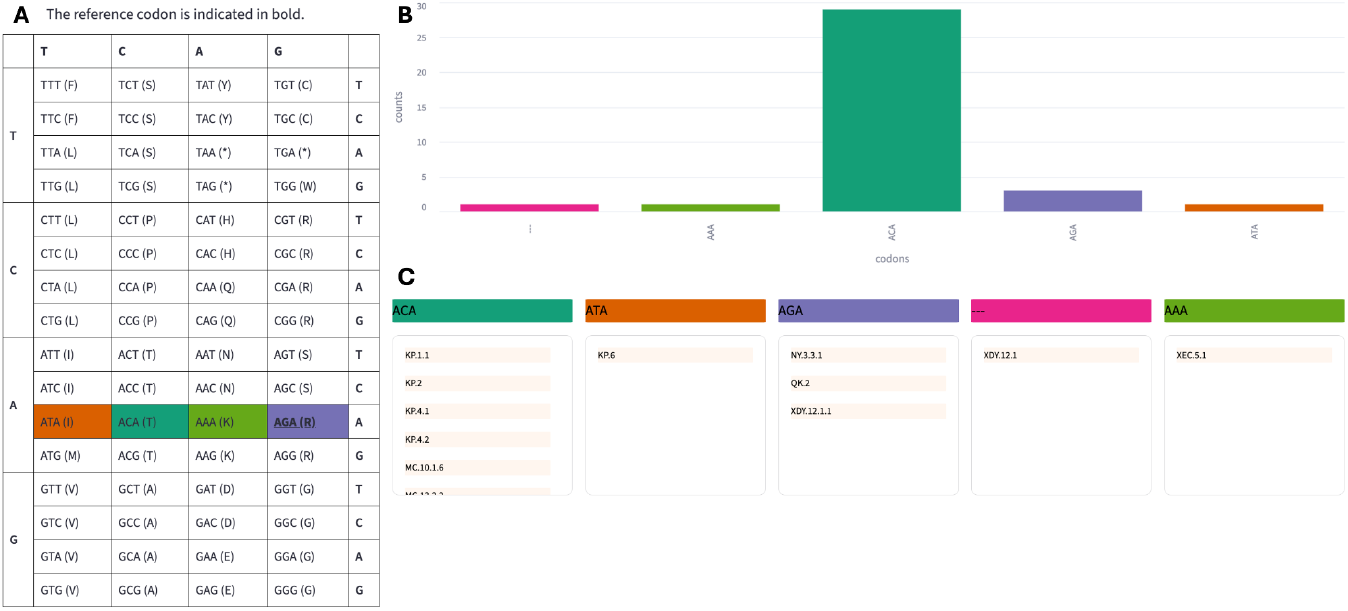
The ConvMut Codon exploration view for the input selection JN.1.11.1 (root) and 346 (position). **(A)** genetic code, **(B**) counts of mutations, **(C)** lineages.

Panel A shows the genetic code (correspondences between nucleotide triplets and the corresponding amino acid residues) where we underline the codon exhibited in the wild type genome in the position of interest *p* and color-highlight the codon present in that position in those lineages that feature an amino acid change in *p* and their parent-delta. Panel B shows the counts of mutations per type of codon, i.e., the number of lineages exhibiting that codon in their parent-delta, in the selected position. Panel C details the lineages contributing to those counts.

Finally, Figure 5 provides the visualization built within the *Mutation exploration* tab for `root = JN.1.11.1` and Spike protein position 346.

**Figure 5.**
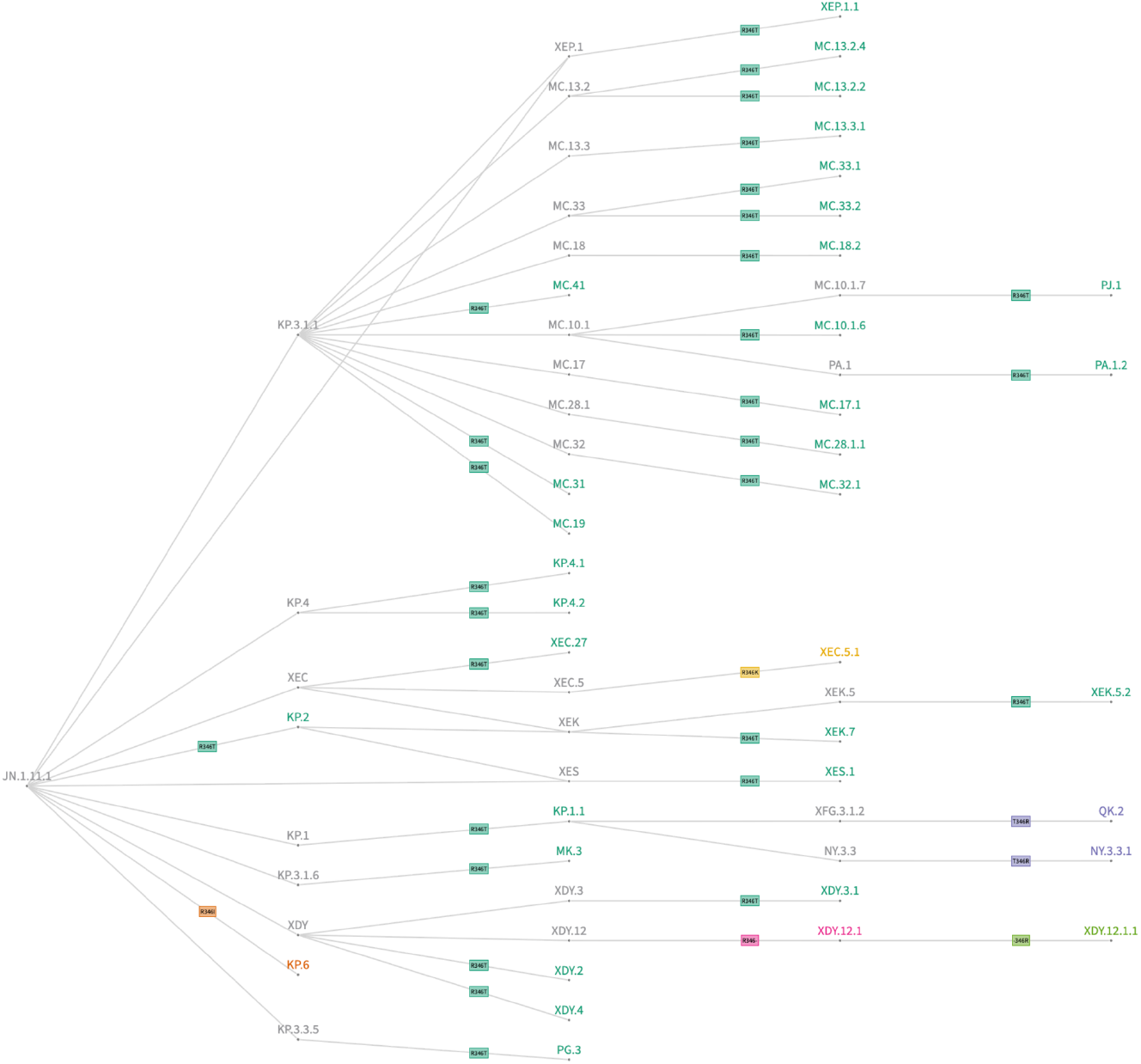
The ConvMut Mutation exploration tree, for root JN.1.11.1 and position 346. Labels on edges indicate the different mutation occurred on this position and acquired by the node on the right of the edge. Note that we use different colors to mark different exhibited amino acid residues.

## Discussion

The analysis of convergent evolution in a pathogen is of paramount importance to predict the changing landscape where antiviral drugs have to act. Clearly, convergent evolution is more evident for protein antigens that are targeted by the selective pressure of the immune response, either infection- or vaccine-elicited. Throughout the four years of the COVID-19 pandemic, the clearest example of convergent evolution has been within the Spike receptor-binding domain (RBD): it has happened in waves and has led to dramatic failures of recently authorized therapeutic anti-Spike monoclonal antibodies [17]. While most of the effort has historically been on studying the divergence in order to provide granular phylogenetic trees, the myriad of sublineages identified for SARS-CoV-2 have posed challenges for the classification of variants. Scientists in the community have volunteered to keep track of shared sets of Spike mutations regardless of the origin and to represent charts that could be easily interpreted by both the general public and vaccine/antibody manufacturers.

Our study has several limitations. First, ConvMut is dependent on the PANGO nomenclature [8], whose designation criteria are somewhat arbitrary and not yet agnostic. While ConvMut is agnostic to support different input phylogenies, currently, the only complete and publicly accessible, up-to-date data source is the one embedded and fed through GISAID pipelines. Second, for recombinant lineages, ConvMut is not able to discriminate the parental source of a given convergent mutation, but -in many instances- the exact breakpoint has not been discriminated against. Third, when estimating the commonest convergent mutations, ignoring the intermediate PANGO-undesignated nodes could have led to an overestimate of mutation counts.

The strengths of ConvMut are: 1) the uniqueness of each of its tools among the plethora of SARS-CoV-2 bioinformatics tools generated so far; 2) the automated updates to GISAID, which is currently the most timely data source for SARS-CoV-2 genomes, lineage assignment and counts; 3) the availability as a user-friendly graphical interface; and 4) the possibility to export the generated charts.

ConvMut can prove useful at forecasting trends in SARS-CoV-2 evolution, benefiting not just basic virological research, but also drug and vaccine manufacturers: the latter can, in fact, adjust their pipeline according to trends, minimizing failures (and their associated economic costs) and focus on drugs that have a higher likelihood of market persistence. This is increasingly important in light of the recent developments in immunobridging [18], which can shorten the time from drug design to market authorization. The architecture of ConvMut is prone to replication on viral pathogens other than SARS-CoV-2 whenever granular phylogenies have been generated.

## Conclusion

We have here exposed the rationale behind and the generation of a novel Web tool to automatically explore convergent evolution in SARS-CoV-2 proteins. Our model could be readapted to different human pathogens for which granular phylogenies are available. Embedding the ConvMut tool within the GISAID framework provides a completely up-to-date instance at the service of immunologists and practitioners around the world, contributing to designing strategies to combat the disease.

## Declarations

### Data Availability Statement

The data employed in ConvMut is made available by GISAID to all users with platform registration.

### Code Availability Statement

The ConvMut software is usable on GISAID for all users with platform registration.

## Acknowledgments

We gratefully acknowledge all data contributors, i.e., the Authors and their Originating laboratories responsible for obtaining the specimens, and their Submitting laboratories for generating the genetic sequence and metadata and sharing via the GISAID Initiative, on which this research is based. We are thankful to the GISAID science and developers teams for building the constellations data generation pipeline and for supporting the whole process of ConvMut as an application within the GISAID platform.

## Institutional Review Board Statement

*Not applicable*.

## Informed Consent Statement

*Not applicable*.

## Author contributions statement

T.A. led the implementation of the application; A.B. supervised the development of the platform and wrote the initial draft of the manuscript; E.F. implemented the first prototype of the application; T.A., A.B., and E.F. jointly designed the computational methods and the Web interface; C.G. and F.M. validated the methods and revised the manuscript. D.F. generated the project and revised the manuscript.

